# Zero is not absence: censoring-based differential abundance analysis for microbiome data

**DOI:** 10.1101/2023.07.05.547842

**Authors:** Lap Sum Chan, Gen Li

## Abstract

Microbiome data analysis faces the challenge of sparsity, with many entries recorded as zeros. In differential abundance analysis, the presence of excessive zeros in data violates distributional assumptions and creates ties, leading to an increased risk of type I errors and reduced statistical power. To address this, we developed a novel normalization method, called CAMP, for microbiome data by treating zeros as censored observations, transforming raw read counts into tie-free time-to-event-like data. This enables the use of survival analysis techniques, like the Cox proportional hazards model, for differential abundance analysis. Extensive simulations demonstrate that CAMP achieves proper type I error control and high power. Applying CAMP to a human gut microbiome dataset, we identify 60 new differentially abundant taxa across geographic locations, showcasing its usefulness. CAMP overcomes sparsity challenges, enabling improved statistical analysis and providing valuable insights into microbiome data in various contexts.

## Part I

## Main Text

### 1 Introduction

Differential abundance analysis is essential for unraveling hidden links between disease and microbiome composition. The goal of differential abundance analysis is to identify taxa that are associated with changes in disease or experimental conditions [1]. Differentially abundant taxa provide insight into how the experimental condition affects the microbiome or how changes in the microbiome are related to disease pathology.

However, performing differential abundance analysis is challenging due to the unique characteristics of microbiome data. Microbiome data often relies on sequencing procedures that primarily yield relative abundances. An essential aspect of such data is compositionality, where the sequencing results provide relative insights into the microbiome profile while being constrained by a unit sum or constant sum restriction. Another prominent feature is that the data are often highly sparse, with excessive zero counts that could be as high as 90% [2].

In the literature, two main approaches to conducting differential abundance analysis are reference-based and proportion-based. These two approaches revolve around testing distinct hypotheses. Reference-based methods evaluate the disparity in absolute abundance, whereas proportion-based methods focus on assessing the observed difference in relative abundance. Each approach has its assumptions and advantages, and the selection between them may hinge upon the particular context of the analysis. In particular, reference-based methods are usually based on parametric or non-parametric tests of log-ratio-transformed data (e.g., ANCOM [3, 4], ANCOM-BC [5], DR [6], and ALDEx2 [7]). The commonly used log-ratio transformations (e.g., centered log-ratio or CLR) lift the compositional data from a simplex into the Euclidean space, allowing standard statistical methods, such as linear models, to be used. However, it is important to note that all reference-based methods rely on the choice of an appropriate reference and assume that this reference is non-differentially abundant, meaning it serves as the baseline or null condition for comparison. Selecting the reference is challenging, and if it is not truly null, observed differences may be artifacts rather than biological distinctions, leading to potential bias in the results.

Proportion-based methods (or count-based methods), on the other hand, provide an alternative perspective that does not require selecting a reference. Commonly used proportion-based methods include corncob [8], LDM [9], metagenomeSeq [2], and ZINQ [1], as well as generic differential analysis methods designed for RNA-Seq data such as DESeq2 [10]. These methods concentrate solely on comparing proportions (or counts) between conditions on a per-taxon basis, eliminating the need to assess the ratios of proportions. Under the assumption that relative abundances hold significant biological implications, proportion-based methods have the ability to identify taxa exhibiting differential relative abundances.

Existing proportion-based methods often encounter challenges such as low statistical power and inflated type-I error rates. These issues are primarily attributed to excessive zeros in microbiome data. The distributional assumptions required by parametric approaches are often violated, and nonparametric tests face additional obstacles posed by tied values resulting from multiple zeros.

To address the issues induced by zeros, we build a new paradigm that treats zeros as censored observations in microbiome data. We develop a novel framework called **c**ensoring-based **a**nalysis of **m**icrobiome **p**roportions (CAMP) that converts zero-inflated microbiome read counts into time-to-event-like data without ties. Consequently, existing survival analysis methods, such as the log-rank test and the Cox proportional hazards models can be leveraged for differential abundance analysis. We demonstrate through extensive simulations that CAMP maintains type I error control and achieves high statistical power. Additionally, using the Cox proportional hazards model, our CAMP approach offers the advantage of enabling adjustment for additional covariates. We applied CAMP to analyze human gut microbiota data from Yatsunenko et al. [11] and identified 60 additional differentially abundant taxa across three pairs of countries compared to existing proportion-based methods.

### 2 Results

#### 2.1 Toy example

At the population level, we begin with a toy example (see Figure 1) to illustrate the difference between reference-based and proportion-based methods. Suppose we have six taxa in two populations and are interested in identifying taxa with differential (relative) abundances. The taxa represented by the blue circle, red pentagon, and orange square have the same relative abundances between populations.

**Fig. 1.**
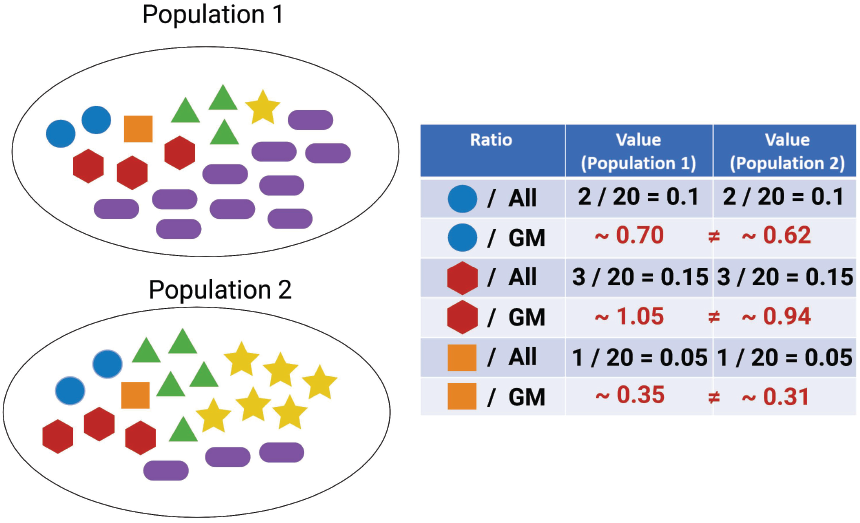
Toy example displaying two populations, each composed of 6 distinct microbial taxa. Each population has a total volume of 20 microbes, with each taxon represented by a unique shape and color. As shown in the table, the taxa of primary interest are the blue circle with a relative abundance of 2/20 = 0.1, the red pentagon with a relative abundance of 3/20 = 0.15, and the orange square with a relative abundance of 1/20 = 0.05 in both populations. The table also provides the ratio of the abundance of each of these three taxa compared to the geometric mean abundance of all taxa within their respective populations. Notice that none of the calculated ratios are identical across the two populations. GM: geometric mean; All: all taxa (blue circle + red pentagon + orange square + green triangle + yellow star + purple ellipse).

In contrast, the taxa symbolized by the green triangle, yellow star, and purple ellipse show differential abundance. More specifically, the green triangle has relative abundances of 3/20 = 0.15 in population 1 and 5/20 = 0.25 in population 2. The yellow star has relative abundances of 1/20 = 0.05 in population 1 and 6/20 = 0.3 in population 2. Lastly, the purple ellipse shows relative abundances of 10/20 = 0.5 in population 1 and 3/20 = 0.15 in population 2.

For reference-based methods, we must pick a reference and determine whether the ratios between a taxon and the reference differ between populations. In particular, if we use the geometric mean (GM) of the relative abundances in a population as the reference, we will encounter spurious discoveries. Specifically, since the GMs of the two populations are distinct, the ratios of the blue circle, red pentagon, and orange square taxa to the GM are different between both populations. Consequently, we might incorrectly conclude that the blue circle, red pentagon, and orange square taxa are differentially abundant, despite their non-differential relative abundances.

The toy example demonstrates that reference-based methods can lead to false positives and potentially invalid results if the reference is misspecified. In general, the accuracy and validity of the results obtained from reference-based methods greatly depend on the selection of a suitable reference that aligns with the assumptions of non-differential abundance. Given that such assumptions are unverifiable in practice and that we typically do not observe absolute abundance, we shift our focus toward relative abundance testing. Such testing does not rely on a reference and aims to assess differential abundance solely based on relative abundance data without extrapolating to absolute abundance. The testing results offer a straightforward interpretation and provide an alternative perspective for differential abundance analysis.

#### 2.2 Overview

Real microbiome data consist of excessive zeros. Some are attributed to the absence of a taxon in a sample, while others are due to sampling issues such as insufficient sequencing depth (i.e., the actual count below the detection limit). Regardless of the underlying data-generating mechanism, a unified way to view zero values in sequencing data is to treat them as **partially observed data**. Namely, zeros are left-censored read counts at the prefixed detection limit. In other words, we only know the unobserved “true” read count is below the limit, but we do not know by how much (or whether it is truly zero or not). Unlike existing zero-replacement strategies, the censoring representation accurately characterizes the information in the sequencing data without making any distributional assumption or introducing unwanted bias.

Correspondingly, we propose a novel normalization workflow for microbiome read counts with excessive zeros. This workflow is visually summarized in Figure 2. The input to the workflow comprises read counts for each taxon, which are then subjected to preprocessing (censoring normalization). Specifically, the zeros in each sample are first converted into a censoring representation at a predetermined cutoff. In order to account for heterogeneous library sizes across different samples, the counts are transformed into relative abundances following conventional practice. Following this, a negative logarithmic transformation is applied to the left-censored relative abundance data. This transformation results in non-negative, right-censored values, allowing for further analysis.

**Fig. 2.**
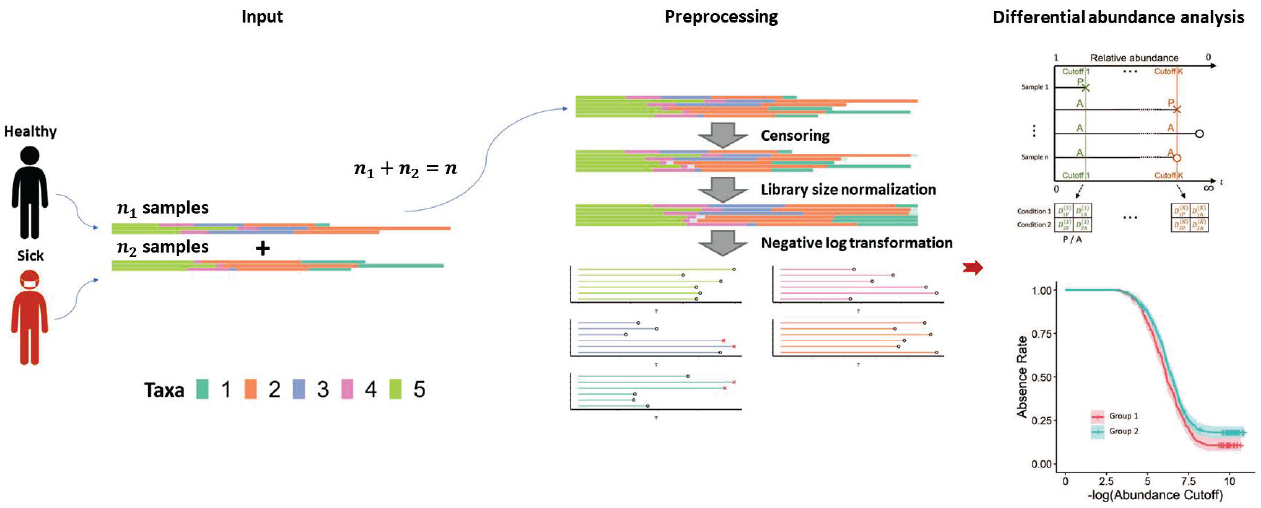
Schematic representation of the CAMP workflow, divided into three key components: input, preprocessing, and differential abundance analysis. The input encompasses read count data from *n* individuals, depicted as bar plots. During preprocessing (by censoring normalization), the read count data is subjected to a three-step process that includes censoring, library size normalization, and a negative log transformation. After censoring normalization, the data from each individual taxon can be represented as censored time-to-event data. The final stage, differential abundance analysis, involves the creation of a presence/absence (P/A) table, computed for the log-rank test, with results visualized via a Kaplan-Meier curve. 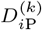 and and 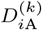: the number of samples present and absent in condition *i* for a given taxon at cutoff *k*, respectively.

Analogous to the right-censored time-to-event data in survival analysis, the transformed microbiome abundance data have an intriguing biological interpretation. In particular, the relative abundance level (from high to low, in a negative log scale) corresponds to the “time” in survival analysis, and the presence of a taxon at a given abundance level is an “event.” Consequently, the concept of time-to-event in survival analysis translates to abundance-to-presence in our transformed microbiome data. The zeros in the raw microbiome data correspond to dropouts or censored observations since they are not “present” at any observed abundance levels. The transformed microbiome data for each taxa can then be visualized using the censoring representation.

Once the microbiome data undergoes censoring normalization, we can leverage existing survival analysis methods to test differential abundance between populations in each taxon. Differential abundance analysis can be performed using a log-rank test. Specifically, we first pool all the uncensored values and define a sequence of abundance cutoffs. At each cutoff, the “at-risk” samples are those with relative abundances at or below the cutoff (or equivalently, with transformed values at or above the observed value). The at-risk samples can be further dichotomized into two mutually exclusive groups based on the presence/absence status of the taxon at the current cutoff, resulting in a *K ×* 2 table (with *K* being the number of populations) and a hypergeometric test statistic. Finally, we aggregate the test statistics across cutoffs to get a single test statistic for the log-rank test, which follows a standard normal distribution under the null after proper normalization. The log-rank test is a nonparametric test that does not impose any distributional assumption on the transformed data and adequately aggregates the presence/absence information across all abundance levels. The transformed values can also be visualized using a Kaplan-Meier curve, facilitating the examination of differences between populations. With covariates, a Cox proportional hazards model can be used, which is elaborated in the Methods section.

#### 2.3 Simulation

We conducted extensive simulations to assess the performance of the proposed CAMP method. In the first simulation, we compared CAMP with five other proportion-based methods (i.e., corncob, DESeq2, LDM, MetagenomeSeq, and ZINQ) to assess their type I error rates and power, using a Dirichlet-multinomial generative model. The sparsity of the data was controlled by varying the multiplicative constant in the Dirichlet distribution, resulting in tables with 30% to 80% zeros. The second simulation setting is similar to the first one, except that we used a truncation approach to induce sparsity in data. The third simulation generated microbiome data from the ZINQ model with corresponding parameters obtained from real data analysis from the CARDIA cohort [12]. In this setting, covariates were also included. We excluded MetagenomeSeq since it cannot handle covariates. Finally, in the fourth simulation setting, we took the real microbiome data from a gut microbiome study [11] and permuted the population labels to evaluate the type I errors of different methods.

In Simulation 1, as shown in panel A of Figure 3, DESeq2 exhibited inflated median type I error rates, ranging from 15% to 50%, as the zero proportion of the data varied from 30% to 80%. Additionally, there was a slight inflation in type I error for corncob, especially when the zero proportion was high. For the other four methods (CAMP, LDM, MetagenomeSeq, and ZINQ), type I error was well-controlled across different zero proportions. In panel B of Figure 3, among the methods with controlled type I error, CAMP achieved the highest power, followed by ZINQ and LDM. Similar trends in type I error rates were found in Simulation 2. In general, as the zero proportion increases, the difference in power between CAMP and other methods becomes bigger, likely due to the fact that CAMP handles zero more rigorously. In panel C of Figure 3, DESeq2 had a baseline type I error rate of 15% at a zero proportion of 30%, which remained relatively constant and reached 22% median type I error when the zero proportion was 80%. On the other hand, corncob became very conservative and reached a 0% type I error when the zero proportion was more than 60%. This resulted in a decrease in power to 0% when the zero proportion was more than 60%, as shown in Panel D of Figure 3. It is interesting to note that LDM and MetagenomeSeq outperformed ZINQ in power in this setting. Nevertheless, the proposed method CAMP still achieved the highest median power while keeping the type I error under control.

**Fig. 3.**
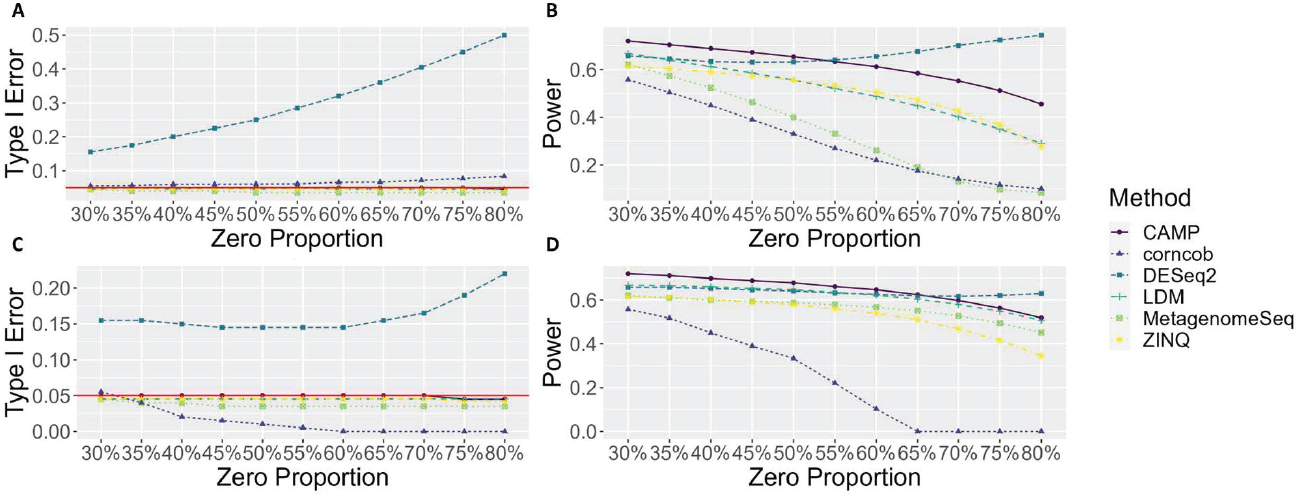
Median type I error and power for 5 methods compared in simulation 1 and 2 across 1000 replicates: (A) type I error for simulation 1; (B) power for simulation 1; (C) type I error for simulation 2; (D) power for simulation 2. Red horizontal line in panel A and C indicates 5% type I error control.

In Simulation 3, the data was generated using ZINQ as the true model. Panel A of Figure 4 shows that CAMP maintained type I error control with a median type I error of 5%, similar to LDM and ZINQ. Conversely, DESeq2, had a median type I error of 22%, and Corncob had 11% median type I error. In terms of power, as shown in panel B of Figure 4, ZINQ achieved a 43% power compared to CAMP’s 40%. This indicates CAMP’s robustness against variations in the data generation mechanism. Though LDM stood as the second most powerful method in controlling type I error in Simulations 1 and 2, its performance waned in the current simulation. In contrast, our proposed method, CAMP, maintained a comparable performance to the true model ZINQ, highlighting the robustness of our non-parametric method against LDM that leans on the form of a multivariate linear regression.

**Fig. 4.**
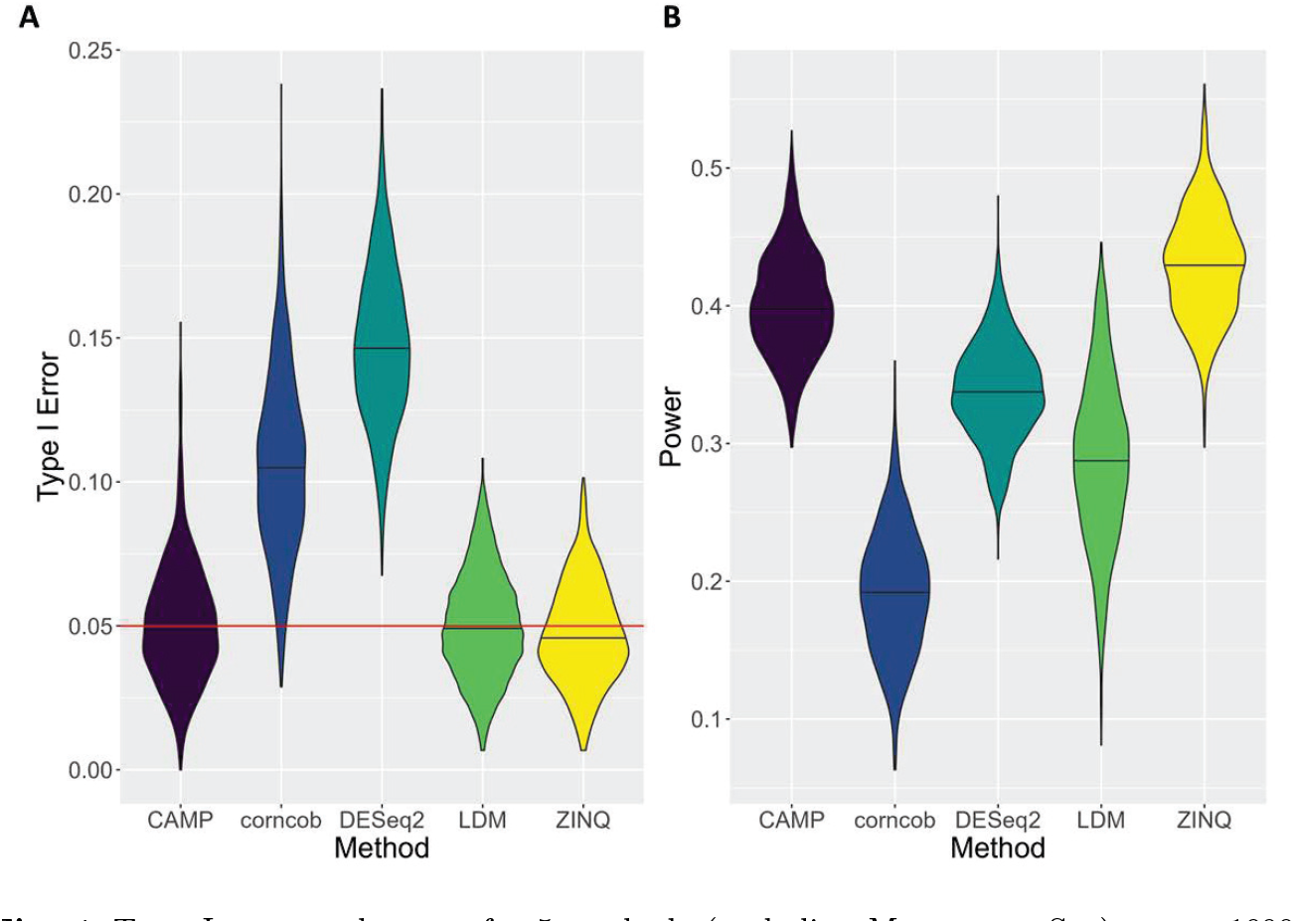
Type I error and power for 5 methods (excluding MetagenomeSeq) across 1000 replicates: (A) type I error for simulation 3; (B) power for simulation 3. Red horizontal line in panel A indicates 5% type I error control.

In Simulation 4 (Figure 5), which utilized real microbiome data with permuted labels, CAMP, LDM, and MetagenomeSeq were the only methods that consistently controlled the median type I error rate. MetagenomeSeq, however, was slightly more conservative than the other two. The remaining methods^−^corncob, DESeq2, and ZINQ^−^exhibited inflated median type I errors, specifically 20%, 12%, and 8%, respectively, across three pairwise comparisons. Once again, this showcases the advantages of employing a non-parametric approach in differential abundance analysis.

**Fig. 5.**
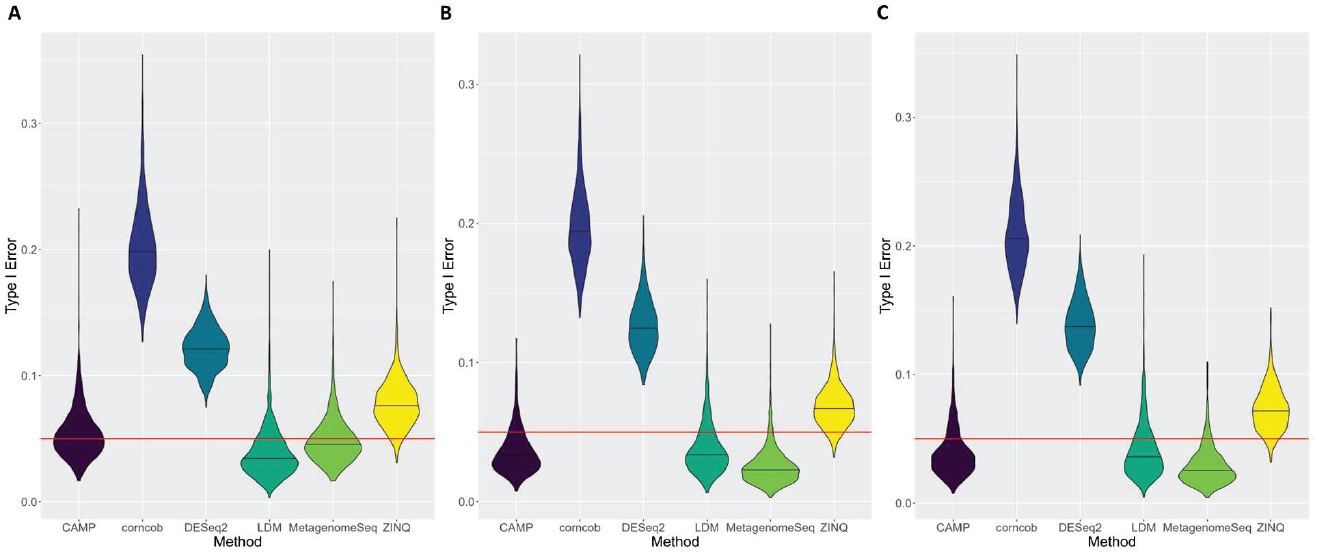
Type I error for 5 methods across 1000 replicates based on pairwise country comparison in simulation 4: (A) US vs Malawi; (B) Malawi vs Venezuela; (C): US vs Venezuela.

#### 2.4 Real data application

We applied the same six proportion-based methods to the human gut microbiome dataset [11] for differential abundance analysis. As an illustration, we present the results of the comparison between Malawi and Venezuela, while the results of the other two comparisons (Malawi vs US and US vs Venezuela) are available in the supplementary materials. When presenting discoveries, we excluded corncob and DESeq2 from the Venn diagram since they had highly inflated type-I errors as shown in Figure 5.

As shown in panel A of Figure 6, 30 taxa were identified to be differentially abundant by all four methods. The proposed CAMP method had a decent amount of overlapping findings with every alternative method (i.e., 84 = 10 + 6 + 30 + 38 with LDM; 42 = 6 + 30 + 1 + 5 with MetagenomeSeq; 100 = 30+38+5+27 with ZINQ). On the other hand, CAMP also had a large number of unique discoveries (i.e., 30), even more than the other methods combined (i.e., 20 = 4 + 1 + 3 + 6 + 1 + 5). Our simulations consistently demonstrated that CAMP effectively controlled the type I error, instilling confidence in these supplementary findings. Further exploration of these distinctive discoveries may yield novel insights into the impact of geography on the gut microbiome. Panels B and C of Figure 6 further provided insights into why CAMP can yield more discoveries. In particular, take Raoultella, a unique discovery in CAMP (*p*-value = 1.8 *×* 10^*−*6^), as an example. At first glance, the violin plot in Panel B showed that the relative abundance distributions of the taxon were similar in the two countries. Upon closer examination of the tail distribution using the proposed censoring-based normalization, the Kaplan-Meier curves revealed distinct separation in the low abundance levels. CAMP, by accurately modeling zeros, demonstrated the ability to detect subtle differences that remain unidentified by other proportion-based methods.

**Fig. 6.**
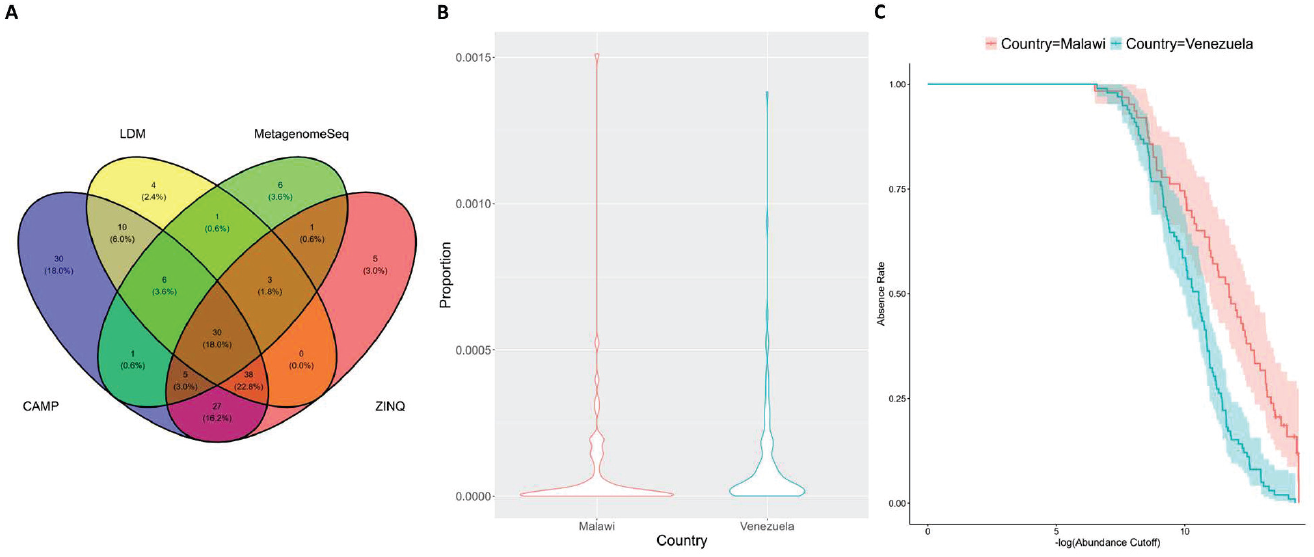
Differential abundance analysis results for gut microbiome dataset: (A) number of discoveries given by 4 methods (excluding corncob and DESeq2) in the Malawi vs Venezuela comparison; (B) proportion distribution for the Raoultella genus in Malawi and Venezuela; (C) Kaplan-meier curve for the Raoultella genus comparing Malawi vs Venezuela.

In total, CAMP yielded 60 unique discoveries across all three pairwise comparisons, which are listed in the Appendix.

### 3 Discussion

We developed CAMP, a censoring-based approach for microbiome data normalization and differential abundance analysis. By transforming zero-inflated read counts to time-to-event-like data, we opened up the possibility of analyzing microbiome data with survival analysis methods. Specifically, we leveraged the log-rank test and Cox proportional hazards model for differential abundance analysis which yields a good power and type I error control.

We envision CAMP is not limited to analyzing microbiome data, but also has potential applications for other omics data types, such as RNA-seq data. While much of the literature considers RNA-seq as count data [13, 14], it is fundamentally compositional [7, 15, 16], as different RNA molecules within a sample sum to a constant. Studies have argued that treating RNA-seq as compositional data is essential [17] and this can yield more robust and reproducible results [7]. As an illustration of how CAMP can be applied to RNA-seq data, we use transcripts per million (TPM), one of the most common normalizations used for RNA-seq [18], as an example. Since the number of transcripts in a sample sum to a million, the data remains compositional after TPM normalization, and CAMP can be directly applied. Even if RNA-seq datasets are already normalized using another common normalization method, RPKM (reads per kilobase of transcript per million reads mapped), the data can still be transformed to TPM [19], thus enabling the potential application of CAMP for differential expression analysis in RNA-seq data. In particular, the application of the CAMP framework can be appealing for microRNA due to its sparsity nature [20].

The CAMP approach can also be generalized to longitudinal differential abundance analysis and differential abundance analysis of multiple taxa using a multivariate frailty model:

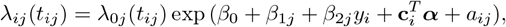

where *j* indexes the longitudinal time point 1, …, *T* or taxa 1, …, *P*, and *a*_*ij*_ is a random effects term known as the frailty. The key distinction between the longitudinal setting and the multiple taxa setting lies in how we model the covariance structure of *a*_*ij*_. For instance, in the longitudinal setting, it might be appropriate to assume *a*_*ij*_ = *a*_*i*_, such that the random effects a cross time points are consistent within the same individual. In the multiple taxa setting, it might be appropriate to model the correlation between random effects across different taxa within the same sample. One possible approach is to assume that the random effects *a* _*ij*_ follow a multivariate normal or multivariate gamma distribution to reflect the correlations between different taxa. This presents a unifying framework in which the current Cox proportional hazards model (without frailty) is a special case. In conclusion, the CAMP framework has great potential to offer a versatile and powerful approach for differential abundance analysis not only in single taxon cross-sectional settings but also in longitudinal and multivariate settings.

While the current CAMP framework has shown promising results and has significant potential applications, there are several areas that could be developed further to expand its usability. One promising avenue for future research could involve devising strategies to handle the inherent compositionality of microbiome data across multiple taxa concurrently, as the existing method primarily focuses on one taxon at a time. Moreover, the extension of the CAMP method to multivariate data sets presents an exciting opportunity for improvement. The existing univariate approach is not immediately applicable to multivariate data. However, the earlier discussions on the potential to use multivariate survival analysis frailty models provide a solid foundation for such advancements. Finally, establishing more flexible and robust strategies for handling detection limits in the CAMP framework could also be an important area for future exploration. Under the current setting, CAMP requires a detection limit for read counts in each sample, and if only compositional data is available, a proxy (like the smallest proportion per sample) needs to be used. This could potentially introduce bias into the analysis, so new strategies to handle this issue would significantly improve the framework’s applicability. By addressing these aspects, the capabilities of the CAMP framework can be significantly enhanced in future iterations.

## Part II

## Methods

### 1 Censoring normalization and connection to survival analysis

In this section, we present a method to handle zero values in compositional microbiome data by treating them as censored observations, censored at the detection limit. Through a series of transformations, we can convert the zero-inflated raw read counts into time-to-event data. Importantly, this transformed data is likely to be without ties, thus addressing the issue of reduced power in nonparametric testing due to ties.

By transforming the microbiome data into a format compatible with survival analysis, we can leverage standard survival analysis techniques for differential abundance analysis. This strategy facilitates more robust and accurate identification of taxa exhibiting differential abundance between conditions, even when faced with zero-inflated data. Below, we elaborate on the concept of censoring normalization, a method designed to censor zeros and convert compositional microbiome data into time-to-event-like data.

Suppose we have *n* samples and *p* taxa. Let **X** = {*x*_*ij*_} be the matrix corresponding to the OTU table of read counts, where *i* = 1, …, *n*, indexes the sample and *j* = 1, …, *p*, indexes the taxa. The library size for sample *i* is given by 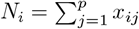.

We define a surrogate read count matrix 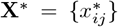 and an indicator matrix **∆** = *{δ*_*ij*_} as follows:

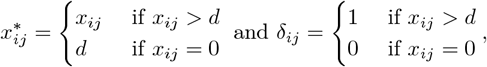

where *d >* 0 is a predefined detection limit in a sequencing study.

The detection limit *d* may vary depending on the context. In scenarios where a predefined detection limit is provided, we directly use this value for *d*. However, there may be instances where a detection limit isn’t explicitly defined. In such cases, it’s necessary to estimate this limit using the available data. In practice, *d* is often the smallest non-zero count (e.g., 1) and can be sample-specific. We can assume the detection limit to be uniform across all samples and taxa, and set 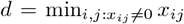. Correspondingly, the new library size for sample *i* is 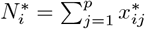.

Left-censoring, in this context, refers to the situation where the read count falls below the detection limit, and we only know that the true value is somewhere below the limit, but not the exact value. Following the convention of survival analysis, if *δ*_*ij*_ = 0, then 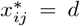 and we can use a shorthand representation 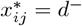, which represents the left-censoring of the read count.

To make the data comparable across samples, we obtain the relative abundance matrix **P** = {*p*_*ij*_} by normalizing the surrogate read counts, i.e., 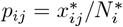. Notably, the matrix **P** no longer contains any zero values, though it may still contain left-censored values. Next, to bridge the left-censored compositional data with the right-censored time-to-event data in survival analysis, we apply a negative log transformation to **P**, resulting in a “time” matrix **T** = *{t*_*ij*_} defined as:

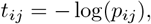

where, by design, *t*_*ij*_ *∈* [0, *∞*) for *i* = 1, …, *n* and *j* = 1, …, *p*.

This transformed matrix, **T**, allows us to draw an analogy with survival analysis. In survival analysis, the notion of censoring is used to handle incomplete information, such as when the event of interest (e.g., death, disease onset) has not occurred by the termination of a study. Applying this idea to microbiome data here, an “event” can be defined as the presence of a taxon at a specific relative abundance level, while the “time” corresponds to the transformed relative abundance values. The “at-risk” population at a given time point consists of samples whose taxa are still absent at this given relative abundance level.

Pictorially, Figure 2 summarizes the key steps in our censoring transformation with *d* = 1 (assumed minimum read counts across all samples).

### 2 Differential abundance analysis

In survival analysis, the time-to-event data is defined by the time of event and the censoring indicator. In our context, through the censoring normalization approach outlined in the previous section, we have converted our microbiome data into time-to-event-like data. Therefore, we are now in a position to apply standard survival analysis techniques for differential abundance analysis.

#### 2.1 Without covariates

Typically, differential abundance analysis is performed for each taxon separately. In this subsection, we will discuss the analysis for a generic taxon. This analysis can be cast as a hypothesis testing problem where the null hypothesis is that the abundance data is identically distributed in different conditions. In scenarios where there is no censoring (i.e., no zero values in the read counts), nonparametric tests such as the Wilcoxon Rank Sum test or the Kruskal-Wallis test can be used to obtain a *p*-value for each taxon. However, in practice, microbiome data are often zero-inflated, resulting in excessive censored values in the transformed data. As a remedy, we propose exploiting the log-rank test [21] in survival analysis, which we will briefly describe below in the context of differential abundance analysis. Without loss of generality, we assume there are only two conditions *k* = 1 and 2 (e.g., healthy vs diseased). Nonetheless, the method can be trivially extended to more than two conditions.

More specifically, we can utilize a “cutoff” value, which refers to a specific threshold value for the transformed relative abundance. This cutoff is used to distinguish the presence or absence of a taxon at that particular abundance level. Consequently, the “at-risk” population at any cutoff *k* is defined as the set of subjects whose transformed data are at or above *k* (equivalently, those whose relative abundances are at or below exp (*−k*)). Among those in this at-risk group, an event is considered to have occured for subjects with transformed data exactly equal to *k* without censoring, meaning that the taxon becomes present in these subjects at the corresponding relative abundance level. For subjects whose transformed data are either above *k* or censored at *k*, they do not have an event, indicating that the taxon remains absent at the given relative abundance level. As such, the at-risk population at *k* is split into two mutually exclusive groups based on the presence/absence (P/A) status of the taxon at the corresponding relative abundance cutoff.

In our context, we can define 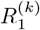 and 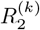 as the at-risk population for conditions 1 and 2 at cutoff *k*, respectively. Then, we define the total at-risk population at cutoff *k* as 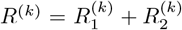. Let 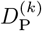 represent the number of events (i.e., presence of a taxon) and 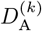 be the count of no events (i.e., absence of a taxon), respectively. It is evident that 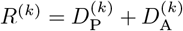.

For a given taxon at cutoff *k*, let 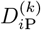 and 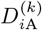 denote the number of samples present and absent in condition *i*. With these counts, we can construct a 2 *×* 2 presence/absence (P/A) table for each at-risk population at cutoff *k* (see Figure 2). Assuming the P/A status of the taxon at cutoff *k* to be independent of the condition under the null, and given the margins of the P/A table 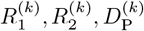 and 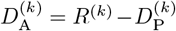 are fixed, the number of events in one condition follows a hypergeometric distribution. For instance, in condition 1, we have the following:

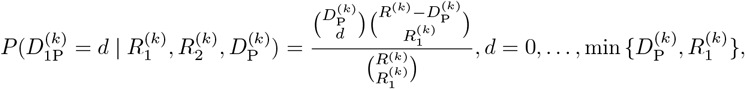

with the mean and variance of 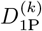 given by

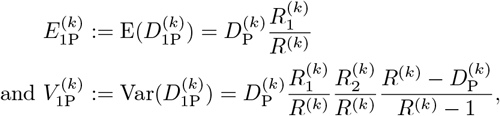

respectively. Following this, the difference between the observed and expected 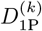 can be computed and summed across different cutoffs to yield a single statistic *U* :

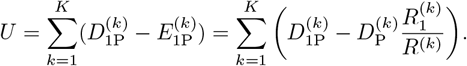

*U* has a corresponding variance *V* :

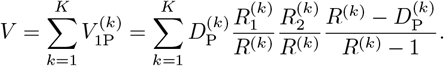

After normalization, the final test statistics, *U/V* ^1/2^ follows the standard normal distribution *N* (0, 1). Alternatively, we could also use *U* ^2^*/V* which follows 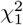. Thus, this enables an explicit *p*-value calculation for the log-rank test.

In this context, the log-rank test adequately aggregates the presence/absence (P/A) information across all abundance levels where an event occurs. At each cutoff, the test statistic evaluates the P/A information to assess the degree of deviation from the null hypothesis. This procedure is done individually at each cutoff, thus capturing the deviation at each level. The final test statistic is an aggregate of these individual deviations, providing a comprehensive summary of the P/A information across all abundance levels. Notice this test is nonparametric that it does not impose any distributional assumption on the data. More importantly, when compared to other nonparametric tests for abundance data that employ ad hoc zero-replacement strategies, our censoring transformation and log-rank test make the most use out of the zero values without additional assumed information.

#### 2.2 With covariates

Sometimes, it is of interest to conduct differential abundance analysis with covariate adjustment. This can be easily achieved by fitting a Cox proportional hazards model:

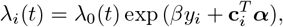

where ***c***_*i*_ *∈ ℝ*^*q*^ denotes a vector of length *q* for covariates and ***α*** represents the corresponding coefficient. In the model above, *λ*_*i*_(*t*) is the hazard function for the *i*th sample for a specific taxon, *λ*_0_(*t*) is the baseline hazard function for the taxon, *y*_*i*_ *∈ {*0, 1} is a binary variable indicating the condition for the *i*th sample, and *β* is a regression coefficient corresponding to the taxon that captures the log hazard ratio between the two conditions. As the hazard function uniquely defines the distribution of a time-to-event random variable, differential abundance analysis is equivalent to testing the hypothesis *H*_0_ : *β* = 0 vs *H*_1_ : *β≠*0, where *β* is the log hazard ratio between the two conditions, while keeping all the covariates constant. In practice, we can first fit the Cox model using the partial likelihood approach. Subsequently, we can use the Wald test, score test, or the likelihood-ratio test (LRT) *−* all of which are asymptotically equivalent tests to test *β* and obtain the *p*-value.

Notice in the absence of covariates, the log-rank test can equivalently be performed using a score test after fitting the Cox proportional hazards model without covariates:

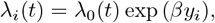

where we test *H*_0_ : *β* = 0 vs *H*_1_ : *β≠*0. In other words, **the Cox model encompasses the log-rank test as a special case**.

Again, it should be noted that this approach is not limited to just two conditions and can easily be generalized to accomodate multiple conditions (e.g., by using multiple dummy variables in the model).

### 3 Simulation

We conducted 4 main simulations to investigate the performance of proportion-based methods. In all 4 simulations listed below, 1000 replicates were used.

#### 3.1 Simulation 1 and 2

In the first two simulations, we considered two groups (conditions), with *n*_1_ and *n*_2_ denoting the number of samples in group 1 and 2, respectively. In both simulations 1 and 2, we set *n*_1_ = *n*_2_ = 100, resulting in a total of *n* = 200 samples and *p* = 1000 taxa. In practice, we do not know the proportion of null taxa. In these settings, we simulate a scenario where a large proportion of taxa is non-null. We set the first 800 taxa to be non-null and the remaining 200 as null taxa.

We generated the initial library size, *N*_*i*_, from Unif(25000, 40000). Sample-specific proportions were generated using Dirichlet(*N*_*i*_, *C***p**_*k*_) where *k* = 1, 2 corresponds to the group index and *C* is a multiplicative constant. We generated all 1000 entries of **p**_1_ from Unif(0, 1), and normalized the first 800 entries to sum up to 0.8. The remaining 200 entries were normalized to have a sum up to 0.2. We followed a similar process for **p**_2_: we first generated the first 800 entries from Unif(0, 1) and normalized them to have a sum of 0.8. The final **p**_2_ vector was constructed by concatenating these 800 normalized entries with the last 200 entries of **p**_1_ (which were normalized to sum up to 0.2). This resulted in two true proportion vectors, **p**_1_ and **p**_2_, with 200 common values (null taxa) and 800 distinct values (non-null taxa).

We generated the Read counts, or the OTU table, using a multinomial distribution with corresponding sample-specific proportions. In simulation 1, we varied the value of *C* to generate different levels of zeros in the OTU table. In simulation 2, we fixed *C* = 353 (approximating 30% zero) and truncated the OTU table at different thresholds, corresponding to our detection limit *d*. This yielded OTU tables with a range of zeros from 30% (at *d* = 0) to 80%.

#### 3.2 Simulation 3

In order to explore further the performance of proportion-based methods, we replicated a simulation from a previous study by Ling et al. [1]. This simulation used the Coronary Artery Risk Development in Young Adults (CARDIA) dataset [12] which includes *n* = 531 samples and *p* = 148 genera. The primary variable of interest in this dataset is the high blood pressure (HBP) status, a binary indicator. Adjustment covariates include age, physical activity, and diet quality score. MetagenomeSeq was excluded from the analysis in this simulation, as it does not allow the inclusion of covariates.

To generate the OTU table, the data was rarefied 10 times to the minimum read depth (46,663), with averages taken to minimize randomness. Covariates were subsequently generated by resampling HBP and the adjustment covariates. The genera were then fitted to the simulated data using the two-part quantile regression model (ZINQ) [1] to obtain parameter estimates. These estimates were then used as the true parameters for the simulation. To assess power, read counts for the 148 genera were regenerated based on the true parameters. To assess type I error, the same process was followed but with the parameter corresponding to HBP set to zero prior to read count regeneration.

#### 3.3 Simulation 4

To complement our previous simulations, which were based on data generated under assumed underlying models, we conducted a simulation using real-world human gut microbiome data from Yatsunenko et al. [11]. More information about this dataset can be found in Section 4 of Methods. After data preprocessing, we performed pairwise differential abundance analysis by comparing the data from two countries at a time and repeating this process for all 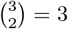 country pairs. To assess the type I error rate, we permuted the country labels for each pair.

### 4 Human gut microbiome dataset and preprocessing

We utilized the human gut microbiome dataset from Yatsunenko et al. [11] in our study. The data was collected from fecal samples of 531 healthy individuals from the Metropolitan areas of United States, Amazonas of Venezuela, and rural Malawi. The dataset was retrieved from the last author’s website https://gordonlab.wustl.edu/supplemental-data/supplemental-data-portal/yatsunenko-et-al-2012/, and the initial OTU table contained 528 samples and 12,251 species. However, due to high sparsity (84.1% zeros) in the initial OTU table, we aggregated the data to the genus level, resulting in *p* = 666 taxa. We also utilized the provided meta data, which included information on the geographical location of each sample. Specifically, there were 63, 315, and 150 samples from Malawi, the United States, and Venezuela, respectively. More details about the dataset can be found in Yatsunenko et al. [11].

### 5 Real data analysis

After preprocessing the human gut microbiome dataset from Yatsunenko et al, we applied 5 proportion-based methods, CAMP, corncob, DESeq2, LDM, MetagenomeSeq and ZINQ to identify differentially abundant taxa across geographical locations. We performed differential abundance analysis for all country pairs by subsetting the data.

## References

[1] Ling, W., Zhao, N., Plantinga, A.M., Launer, L.J., Fodor, A.A., Meyer, K.A., Wu, M.C.: Powerful and robust non-parametric association testing for microbiome data via a zero-inflated quantile approach (zinq). Microbiome 9(1), 1–19 (2021)

[2] Paulson, J.N., Stine, O.C., Bravo, H.C., Pop, M.: Differential abundance analysis for microbial marker-gene surveys. Nature methods 10(12), 1200– 1202 (2013)

[3] Mandal, S., Van Treuren, W., White, R.A., Eggesbø, M., Knight, R., Peddada, S.D.: Analysis of composition of microbiomes: a novel method for studying microbial composition. Microbial ecology in health and disease 26(1), 27663 (2015)

[4] Kaul, A., Mandal, S., Davidov, O., Peddada, S.D.: Analysis of microbiome data in the presence of excess zeros. Frontiers in microbiology 8, 2114 (2017)

[5] Lin, H., Peddada, S.D.: Analysis of compositions of microbiomes with bias correction. Nature communications 11(1), 1–11 (2020)

[6] Morton, J.T., Marotz, C., Washburne, A., Silverman, J., Zaramela, L.S., Edlund, A., Zengler, K., Knight, R.: Establishing microbial composition measurement standards with reference frames. Nature communications 10(1), 1–11 (2019)

[7] Fernandes, A.D., Reid, J.N., Macklaim, J.M., McMurrough, T.A., Edgell, D.R., Gloor, G.B.: Unifying the analysis of high-throughput sequencing datasets: characterizing rna-seq, 16s rrna gene sequencing and selective growth experiments by compositional data analysis. Microbiome 2(1), 1– 13 (2014)

[8] Martin, B.D., Witten, D., Willis, A.D.: Modeling microbial abundances and dysbiosis with beta-binomial regression. The annals of applied statistics 14(1), 94 (2020)

[9] Hu, Y.-J., Satten, G.A.: Testing hypotheses about the microbiome using the linear decomposition model (ldm). Bioinformatics 36(14), 4106–4115 (2020)

[10] Love, M.I., Huber, W., Anders, S.: Moderated estimation of fold change and dispersion for rna-seq data with deseq2. Genome biology 15(12), 1–21 (2014)

[11] Yatsunenko, T., Rey, F.E., Manary, M.J., Trehan, I., Dominguez-Bello, M.G., Contreras, M., Magris, M., Hidalgo, G., Baldassano, R.N., Anokhin, A.P., et al.: Human gut microbiome viewed across age and geography. nature 486(7402), 222–227 (2012)

[12] Friedman, G.D., Cutter, G.R., Donahue, R.P., Hughes, G.H., Hulley, S.B., Jacobs Jr, D.R., Liu, K., Savage, P.J.: Cardia: study design, recruitment, and some characteristics of the examined subjects. Journal of clinical epidemiology 41(11), 1105–1116 (1988)

[13] Kuczynski, J., Lauber, C.L., Walters, W.A., Parfrey, L.W., Clemente, J.C., Gevers, D., Knight, R.: Experimental and analytical tools for studying the human microbiome. Nature Reviews Genetics 13(1), 47–58 (2012)

[14] Anders, S., McCarthy, D.J., Chen, Y., Okoniewski, M., Smyth, G.K., Huber, W., Robinson, M.D.: Count-based differential expression analysis of rna sequencing data using r and bioconductor. Nature protocols 8(9), 1765–1786 (2013)

[15] Quinn, T.P., Richardson, M.F., Lovell, D., Crowley, T.M.: propr: an r-package for identifying proportionally abundant features using compositional data analysis. Scientific reports 7(1), 1–9 (2017)

[16] Quinn, T.P., Erb, I., Richardson, M.F., Crowley, T.M.: Understanding sequencing data as compositions: an outlook and review. Bioinformatics 34(16), 2870–2878 (2018)

[17] McGee, W.A., Pimentel, H., Pachter, L., Wu, J.Y.: Compositional data analysis is necessary for simulating and analyzing rna-seq data. bioRxiv, 564955 (2019)

[18] Abrams, Z.B., Johnson, T.S., Huang, K., Payne, P.R., Coombes, K.: A protocol to evaluate rna sequencing normalization methods. BMC bioinformatics 20(24), 1–7 (2019)

[19] Zhao, S., Ye, Z., Stanton, R.: Misuse of rpkm or tpm normalization when comparing across samples and sequencing protocols. Rna 26(8), 903–909 (2020)

[20] McGeary, S.E., Lin, K.S., Shi, C.Y., Pham, T.M., Bisaria, N., Kelley, G.M., Bartel, D.P.: The biochemical basis of microrna targeting efficacy. Science 366(6472), 1741 (2019)

[21] Mantel, N., et al.: Evaluation of survival data and two new rank order statistics arising in its consideration. Cancer Chemother Rep 50(3), 163– 170 (1966)

