## Appendix for "Zero is not absence: censoring-based differential abundance analysis for microbiome data"

### Part III

### Appendix

#### A Real data analysis results

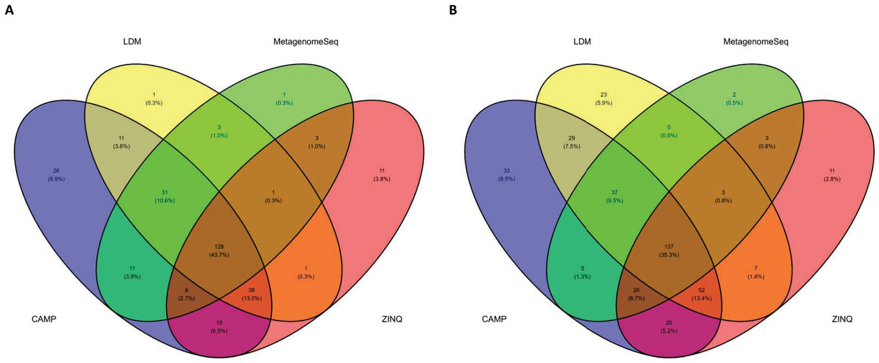


**Fig. 7** Differential abundance analysis results for gut microbiome dataset: (A) number of discoveries given by 4 methods (excluding corncob and DESeq2) in the Malawi vs US comparison; and (B) US vs Venezuela comparison.

**Table 1**: List of 60 taxa (genus) uniquely identified by CAMP in the three pairwise comparisons (MA vs US; MA vs VE and US vs VE) with their corresponding CAMP $p$-values. $p$-values indicating statistical significance (at 5% false discovery rate) are bolded. The maximum significant $p$-values at 5% false discovery rate for MA vs US, MA vs VE, and US vs VE are 0.0191, 0.0110, and 0.0245, respectively. Discoveries unique to CAMP are denoted with an asterisk (*).

| Genus | $p$-value (MA vs US) | $p$-value (MA vs VE) | $p$-value (US vs VE) |
|---|---|---|---|
| A17 | $\mathbf{2.3 \times 10^{-3}}$* | 0.0373 | 1 |
| Actinoallomurus | $\mathbf{2.5 \times 10^{-3}}$ | 0.7016 | **0.0110***|
| Alkalibacterium | 0.5196 | 0.2849 | **0.0112*** |
| Alloiococcus | 1 | 0.1897 | $\mathbf{5.1 \times 10^{-3}}$* |
| Aminiphilus | $\mathbf{9.6 \times 10^{-7}}$* | $\mathbf{9.1 \times 10^{-4}}$* | 1 |
| Amycolatopsis | $\mathbf{2.2 \times 10^{-4}}$ | 0.4241 | $\mathbf{7.1 \times 10^{-3}}$* |
| Aquimonas | **0.0144*** | 0.9142 | $\mathbf{4.6 \times 10^{-3}}$ |
| Arcanobacterium | $\mathbf{5.4 \times 10^{-10}}$ | 0.0146 | $\mathbf{2.2 \times 10^{-3}}$* |
| Arsenicicoccus | $\mathbf{4.9 \times 10^{-5}}$ | 0.3256 | **0.0129*** |

Continued on next page

| Genus | $p$-value (MA vs US) | $p$-value (MA vs VE) | $p$-value (US vs VE) |
|---|---|---|---|
| Asticcacaulis | 1 | 0.2450 | $\mathbf{8.9 \times 10^{-3}}$* |
| Azospirillum | 0.0251 | 0.8319 | **0.0184***|
| Bergeriella | 0.0587 | 0.5748 | $\mathbf{4.7 \times 10^{-3}}$* |
| Brachybacterium | $\mathbf{2.5 \times 10^{-13}}$ | $\mathbf{9.8 \times 10^{-3}}$* | $\mathbf{5.1 \times 10^{-4}}$ |
| Bradyrhizobium | $\mathbf{2.9 \times 10^{-3}}$ | 0.7165 | $\mathbf{6.5 \times 10^{-3}}$* |
| Brenneria | $\mathbf{1.7 \times 10^{-3}}$* | 0.0629 | $\mathbf{1.1 \times 10^{-9}}$ |
| Caldilinea | $\mathbf{2.3 \times 10^{-3}}$* | 0.1196 | 0.0624 |
| Caloramator | **0.0151** | **0.0105*** | 0.5124 |
| CandidatusXiphinematobacter | 1 | 0.3241 | **0.0142*** |
| Cardiobacterium | 0.4580 | 0.2490 | $\mathbf{8.1 \times 10^{-3}}$* |
| Caulobacter | 0.1142 | $\mathbf{1.1 \times 10^{-3}}$* | $\mathbf{1.3 \times 10^{-5}}$ |
| Cedecea | $\mathbf{3.4 \times 10^{-7}}$ | $\mathbf{1.0 \times 10^{-3}}$* | **0** |
| Chelativorans | 1 | 0.2442 | $\mathbf{9.4 \times 10^{-3}}$* |
| Chroococcidiopsis | $\mathbf{8.2 \times 10^{-3}}$* | 0.0136 | $\mathbf{5.8 \times 10^{-11}}$ |
| Cycloclasticus | 1 | 0.1597 | $\mathbf{4.0 \times 10^{-3}}$* |
| Dermabacter | $\mathbf{5.5 \times 10^{-7}}$ | **0.0106*** | 0.2385 |
| Devosia | **0.0160*** | 0.6159 | $\mathbf{1.1 \times 10^{-3}}$ |
| Dokdonella | 1 | 0.2568 | **0.0132*** |
| Dyadobacter | **0.0120*** | 0.2277 | 0.0704 |
| Endozoicimonas | 1 | 0.2444 | $\mathbf{9.6 \times 10^{-3}}$* |
| Escherichia | $\mathbf{3.1 \times 10^{-10}}$ | $\mathbf{9.3 \times 10^{-3}}$* | **0** |
| Ethanoligenens | 0.1995 | $\mathbf{6.2 \times 10^{-3}}$* | $\mathbf{1.2 \times 10^{-3}}$ |
| Exiguobacterium | $\mathbf{1.4 \times 10^{-5}}$ | 0.2445 | **0.0117*** |
| Gallionella | $\mathbf{6.7 \times 10^{-3}}$* | $\mathbf{7.2 \times 10^{-5}}$ | **0** |
| HTCC | $\mathbf{5.7 \times 10^{-4}}$* | 0.7174 | $7.6 \times 10^{-3}$ |
| Herpetosiphon | $\mathbf{2.3 \times 10^{-3}}$* | 0.0374 | 1 |
| Iamia | $\mathbf{2.3 \times 10^{-3}}$* | 0.0374 | 1 |
| Klebsiella | $\mathbf{5.0 \times 10^{-9}}$ | **0.0101*** | **0** |
| Methylophaga | $\mathbf{1.6 \times 10^{-12}}$ | $\mathbf{7.3 \times 10^{-3}}$* | **0** |
| Micrococcus | **0** | $\mathbf{3.7 \times 10^{-7}}$ | **0.0227*** |
| Mitsuaria | **0.0169*** | 0.0178 | 0.7555 |
| Nesterenkonia | **0.0106*** | 0.6073 | $\mathbf{3.2 \times 10^{-3}}$ |
| Nonomuraea | $\mathbf{1.0 \times 10^{-6}}$ | 0.0622 | **0.0206*** |
| Ochrobactrum | 0.4323 | 0.0600 | **0.0203*** |
| Pectinatus | **0.0188*** | $\mathbf{8.7 \times 10^{-3}}$ | 0.2038 |
| Pedomicrobium | 1 | 0.2198 | $\mathbf{5.7 \times 10^{-3}}$* |
| Peptostreptococcus | $\mathbf{3.8 \times 10^{-3}}$* | 0.8274 | $\mathbf{2.9 \times 10^{-3}}$ |
| Planctomyces | $\mathbf{5.4 \times 10^{-3}}$* | 0.0614 | 1 |
| Polynucleobacter | 0.3793 | 0.2619 | $\mathbf{9.5 \times 10^{-3}}$* |
| Prevotella | **0**t | $\mathbf{8.4 \times 10^{-3}}$* | **0** |
| Raoultella | $\mathbf{3.8 \times 10^{-14}}$ | $\mathbf{1.8 \times 10^{-6}}$* | **0** |
| Rheinheimera | $\mathbf{7.1 \times 10^{-3}}$* | 0.0169 | $\mathbf{1.5 \times 10^{-11}}$ |
| Salinibacillus | 1 | 0.2509 | **0.0122*** |

Continued on next page

| Genus | $p$-value (MA vs US) | $p$-value (MA vs VE) | $p$-value (US vs VE) |
|---|---|---|---|
| Salinicoccus | $\mathbf{9.7 \times 10^{-3}}$* | 0.2440 | 0.3244 |
| Sejongia | $\mathbf{2.9 \times 10^{-3}}$ | 0.7112 | **0.0114*** |
| Serratia | $\mathbf{8.0 \times 10^{-13}}$ | $\mathbf{6.6 \times 10^{-4}}$* | **0** |
| Skermanella | 0.0603 | 0.7316 | 0.$\mathbf{7.1 \times 10^{-3}}$* |
| Sporosarcina | 0.0571 | 0.6001 | $\mathbf{7.1 \times 10^{-3}}$* |
| TG5 | 0.7218 | 0.1151 | **0.0185*** |
| Terribacillus | **0.0112*** | $\mathbf{9.9 \times 10^{-3}}$ | $\mathbf{4.0 \times 10^{-13}}$ |
| Xenophilus | $\mathbf{5.4 \times 10^{-3}}$* | 0.5494 | 0.0724 |
